# The addition of FAIMS Increases Targeted Proteomics Sensitivity from FFPE Tumor Biopsies

**DOI:** 10.1101/2022.02.08.479554

**Authors:** Steve Sweet, David Chain, Wen Yu, Philip Martin, Marlon Rebelatto, Andrew Chambers, Fabiola Cecchi, Yeoun Jin Kim

## Abstract

Mass spectrometry-based targeted proteomics allows objective protein quantitation of clinical biomarkers from a single section of formalin-fixed, paraffin-embedded (FFPE) tumor tissue biopsies. We combined high-field asymmetric waveform ion mobility spectrometry (FAIMS) and parallel reaction monitoring (PRM) to increase assay sensitivity. The modular nature of the FAIMS source allowed direct comparison of the performance of FAIMS-PRM to PRM. Limits of quantitation were determined by spiking synthetic peptides into a human spleen matrix. In addition, 20 clinical samples were analyzed using FAIMS-PRM and the quantitation of HER2 was compared with that obtained with the Ventana immunohistochemistry assay. FAIMS-PRM improved the overall signal-to-noise ratio over that from PRM and increased assay sensitivity in FFPE tissue analysis for four (HER2, EGFR, cMET, and KRAS) of five proteins of clinical interest. FAIMS-PRM enabled sensitive quantitation of basal HER2 expression in breast cancer samples classified as HER2 negative by immunohistochemistry. Furthermore, we determined the degree of FAIMS-dependent background reduction and showed that this correlated with an improved lower limit of quantitation with FAIMS. FAIMS-PRM is anticipated to benefit clinical trials in which multiple biomarker questions must be addressed and the availability of tumor biopsy samples is limited.

## INTRODUCTION

In targeted therapeutics, expression levels of primary drug targets are often associated with patients’ clinical responses to treatment and thus serve as predictive biomarkers. Precise measurement of such biomarkers plays a crucial role in selecting patients most likely to benefit from therapy ^1^. The U.S. Food and Drug Administration has approved companion diagnostic (CDx) assays to select eligible patients for treatment on the basis of molecular characteristics of the tumor ^2^. To date, CDx assays have been approved for just five protein targets, namely, ALK, EGFR, HER2, KIT, and PD-L1, and all of these protein assays are based on immunohistochemistry (IHC) ^3^. For example, the HercepTest was the first protein-based CDx assay that identified HER2-positive cancer patients eligible for trastuzumab treatment ^4^. Currently, an IHC stain intensity of 3+, as annotated by a pathologist, is considered HER2 positive. A HER2 IHC of 2+ is also considered HER2 positive if an additional fluorescence in situ hybridization assay (FISH) proves that the gene is amplified. Although clinical studies have demonstrated a strong correlation between IHC-based HER2 expression level and trastuzumab sensitivity, a series of mixed results from recent clinical trials ^5^ and ambiguous quantitation of the antibody-based HercepTest necessitates more objective and quantitative protein assays ^6^. Furthermore, recent advances in antibody-drug conjugate (ADC) therapies that conjugate a small-molecule drug to trastuzumab demonstrate that such ADCs can efficiently target tumor cells with low HER2 expression by leveraging the conjugated warheads in heterogeneous tumors ^7^. This new focus on tumors with low HER2 expression highlights an important need for more sensitive and quantitative protein assays.

In addition to patient stratification, quantitative analysis of protein biomarkers in biopsy samples is commonly applied to assess the pharmacodynamic effects of drugs under clinical investigation. This includes monitoring the abundance of therapeutic target proteins as well as downstream efficacy markers upon drug treatment ^8, 9^. IHC is also a standard method for pharmacodynamic assessments in clinical trials.

Mass spectrometry (MS)-based targeted proteomics with selected reaction monitoring (SRM) has emerged as a promising technology for protein quantitation to overcome the limitations of IHC. In reports of clinical trials using SRM-based HER2 measurements, clinical cutoff values of HER2 expression level, which is predictive of response to trastuzumab, were determined by SRM readouts ^6, 10^. The SRM method, using a triple-quadrupole mass spectrometer, is known for its high sensitivity, which is achieved by fixing the mass analyzer to transmit only selected ions. However, sensitivity can be hampered by high non-specific background signal that interferes with the signals of targeted ions. This is a very common problem in analyzing high-complexity clinical samples and has been addressed by using parallel reaction monitoring (PRM) to leverage high-resolving power mass analyzers, allowing a flexible choice of fragment ions postacquisition ^11–14^. With greater signal-to-noise ratios than SRM, PRM is advantageous when an assay suffers from high levels of non-specific background signal, as is often the case for clinical analyses involving formalin-fixed, paraffin-embedded (FFPE) tumor biopsies ^15, 16^.

In theory, additional reduction of unwanted signals may further improve assay performance. Although sample fractionation or enrichment strategies can be applied to reduce complexity and non-specific background signals ^17, 18^, they also decrease the overall throughput and, more importantly, are not feasible when there is limited availability of clinical samples. Ion mobility spectrometry (IMS) can reduce complexity in the gas phase without having to fractionate samples in advance and has been applied to peptide quantitation by SRM ^19–22^.

High-field asymmetric waveform IMS (FAIMS) is a commercially available technology that is distinct from other IMS techniques. In FAIMS, ions are transported by a flow of carrier gas through an asymmetrical electric field applied perpendicularly to the direction of travel ^23^. Separation therefore results in a continuous flow of a subset of ions that are selected by a DC compensation voltage (CV), as opposed to the temporal separation of ions that characterizes other IMS approaches. This feature may be ideal for a targeted proteomics workflow, in which the focus is on a selected list of molecules. In this context, the addition of FAIMS may further increase the performance of PRM. To date, the application of FAIMS on an Orbitrap instrument for PRM has been described in a single publication, without extensive characterization of the benefit of FAIMS ^24^. Here we present a systematic evaluation of a FAIMS-PRM method combining ion mobility and a high-resolution Orbitrap mass analyzer to achieve high-sensitivity quantitation of critical biomarkers in oncology clinical studies.

## MATERIALS AND METHODS

### Synthetic peptide preparation

All peptides were synthesized as light and heavy pairs by 21st Century Biochemicals (Marlborough, MA). Heavy peptides were labeled with C-terminal R [^13^C_6_^15^N_4_] or K [^13^C_6_^15^N_2_] with >99% isotopic enrichment. For cysteines, carbamidomethylated cysteines were used. Synthesized peptides were further purified by HPLC to attain >95% purity, and net peptide content was verified by amino acid analysis.

### Clinical sample preparation

FFPE tumor biopsy samples from breast cancer patients, collected under Institutional Review Board approval by certified medical pathologists, were purchased from ProteoGenex (Inglewood, CA). Three tissue sections per sample were generated for hematoxylin and eosin (H&E) staining, HER2 IHC, and laser microdissection (LMD).

### Image analysis and LMD

Pathology evaluation for laser-microdissected tumor epithelium was conducted by the study pathologist on H&E-stained slides, which were digitally imaged with an Aperio ScanScope AT scanner (Leica Microsystems, Wetzlar, Germany). HALO AI (Indica Labs, Albuquerque, NM) was used to classify and annotate the slides to guide LMD of tumor epithelium (Supplemental Materials and Methods).

### Sample processing for LC-MS analysis

Microdissected tissue samples collected from the slides were dissolved in 0.1% RapiGest, incubated at 95°C for 90 min, alkylated with chloroacetamide at 37°C, prior to overnight trypsin digestion (Supplemental Materials and Methods).

### LC-MS data acquisition

Desalted samples (1.2 μg) were combined with synthetic isotope-labelled peptides (6 fmol). Five-sixths of this mixture (1 μg of total peptide and 5 fmol of each synthetic peptide) was loaded onto EvoTip trapping columns before separation with the EvoSep One nanoLC system (EvoSep, Odense, Denmark) coupled to an Orbitrap Fusion Lumos mass spectrometer with a FAIMS-PRO interface (Thermo Fisher). Peptides were eluted over a 44-min gradient, from 7% to 30% acetonitrile (on-column), at a flow rate of 500 nL/min (Supplemental Materials and Methods). The FAIMS-PRM experiment employed higher-energy collisional dissociation (HCD) fragmentation with an isolation window of 0.7 mass-to-charge ratio (m/z), a target automatic gain control of 1E6 ions, and a maximum injection time of 100 ms. Tandem MS (MS/MS) scans were acquired in centroid mode with the Orbitrap detector, using 30K resolution at 200 m/z (unless otherwise stated it the text). FAIMS was operated at the standard resolution, with no additional FAIMS gas.

### Data analysis

PRM data was analyzed with Skyline (University of Washington, Seattle, WA) ^25^, using high-selectivity extraction. Fragment ions with interference were identified by manual analysis, comparing coelution and fragment ion ratios between endogenous and reference peptides. Any fragment ions showing interference were flagged and omitted from use in quantitation (Supplemental Materials and Methods).

Ion injection numbers were extracted from raw files as RawOvFtT, using rawDiag ^26^.

### HER2 IHC assay

FFPE tissue sections were stained with the Ventana Pathway anti-Her2/neu (4B5) assay according to the manufacturer’s instructions on a Ventana Benchmark Ultra staining device (Ventana Medical, Oro Valley, AZ).

## RESULTS

### Integration of FAIMS into the PRM method

In this study, we selected 10 peptides representing HER2, EGFR, ER, cMET, and KRAS, which are primary targets of oncology drug development and actionable biomarkers in the clinic ^5, 8, 9, 27^. To determine the optimal FAIMS CV that transmits the maximum ion flux, a range of CVs from 0 to –100 V was scanned per precursor, with direct infusion of synthetic peptides. For example, the optimal transmission of the EGFR IPLENLQIIR [M+2H]2+ precursor was between –62 and –58 V, with a half-maximal intensity of –72 to –52 V (Figure 1A). The optimal CV value determined per precursor was integrated into the final scheduled PRM method. This allowed the FAIMS unit to apply the optimal CV values for the targets at the scheduled acquisition time of the individual peptide, according to the PRM method. The final FAIMS-PRM method for these peptides is summarized in Table 1. FAIMS CVs for the 10 peptides ranged from –28 to –58 V. For each target the optimal collision-induced dissociation energy was also determined (Table 1). Additional optimization of the precursor isolation width and MS/MS resolution was carried out. Precursor isolation widths of 0.4, 0.7 and 1.6 m/z were compared: 0.7 m/z was selected as the preferred balance between reducing interference, shown in Supplementary Figure S1A for the HER2 ELVSEFSR peptide, and maintaining signal of the target peptide (Supp. Fig S1B). MS/MS resolutions of 30K (64 ms transient length), 60K (128 ms) and 120K (256 ms) were compared: the additional scan time required for higher resolution data acquisition did not appear to give sufficient reduction in interference to be warranted. Higher resolution acquisition also resulted in lower fragment ion signal (Supp. Fig. S2), and therefore the 30K resolution was preferred. For the four peptides shown in Supplementary Figure S2, the difference in summed area between the 30K resolution and 120K resolution ranges from 1% to 29%, with the median percentage drop being 22%. The lower signal with higher resolution acquisition may be related to the longer measurement time in the Orbitrap analyzer, allowing signal decay to occur, for instance due to collisions with background gas ^28^.

**Figure 1.**
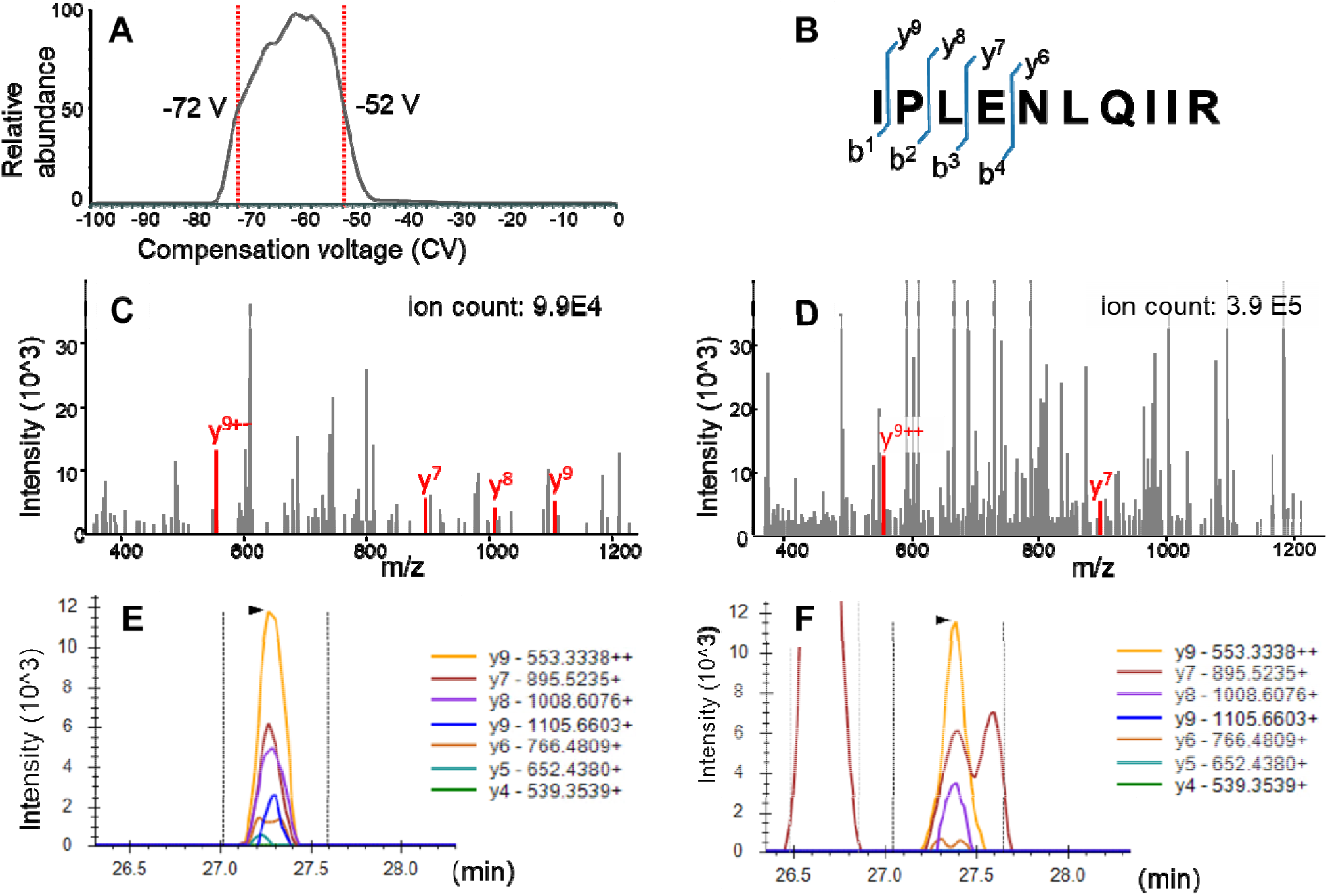
FAIMS effect on PRM analysis of EGFR IPLENLQIIR [M+2H]^2+^. (A) CV scan for EGFR IPLENLQIIR [M+2H]^2+^ with direct infusion of synthetic peptide, using FAIMS CVs from −100 to 0 V, step size 1. The plot shows the MS/MS intensity of the precursor (604.87 m/z) with HCD collision energy set to 0. CVs corresponding to half-maximal intensity are labeled. (B) Fragmentation map of IPLENLQIIR. (C, D) MS/MS spectrum at the apex of 46-amol EGFR IPLENLQIIR elution spiked into trypsin-digested, formalin-fixed spleen (C) with the optimized FAIMS CV and (D) without FAIMS. Fragment ions derived from EGFR IPLENLQIIR are labeled and colored. Ion count refers to total ion count in the Orbitrap cell, recorded as RawOvFtT in the scan header. (E, F) XICs of PRM fragment ions of IPLENLQIIR [M+2H]^2+^ at 46 amol/μg with (E) FAIMS-PRM and (F) PRM.

**Table 1.**
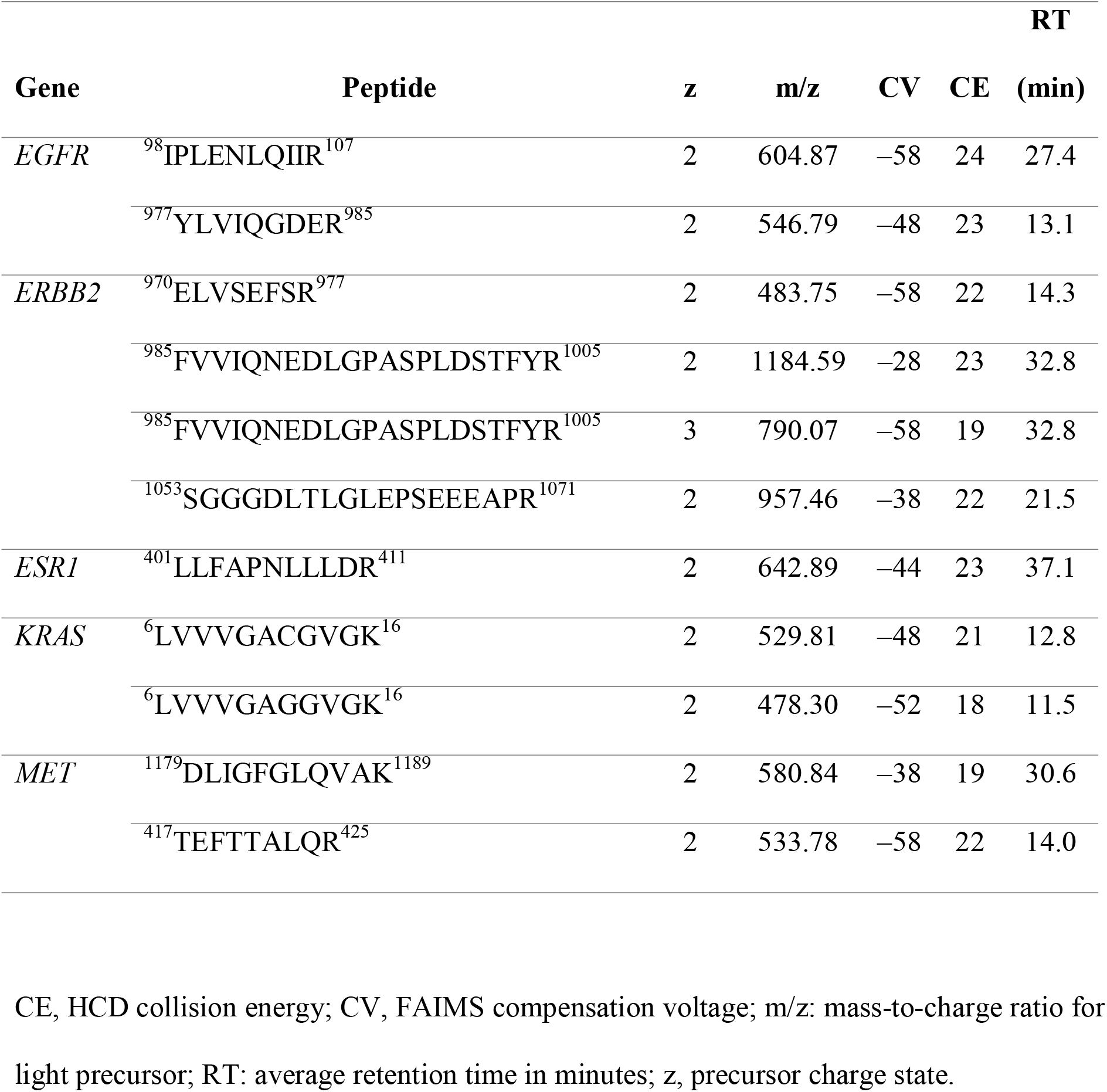
FAIMS-PRM method

### Performance of FAIMS-PRM

To assess the impact of FAIMS integration into the PRM method, the limits of quantitation (LOQs) of the PRM assays obtained with and without FAIMS were determined in trypsin-digested, formalin-fixed human spleen. Because the target proteins were present in the spleen matrix, a reverse-dilution approach was chosen, with spiking-in of a constant 5 fmol of light peptide and a 10-point dilution series of heavy peptide ranging from 100 fmol to 5 amol. Blank (light peptides plus matrix) and double-blank (matrix only) samples were included in this set.

An LOQ curve was acquired on two separate occasions, with and without FAIMS, for a total of four LOQ curves and 192 raw files. The quantitative range, spanning from lower to upper limits of quantitation (LLOQ to ULOQ), was defined as the points on the curve with CV values of <20%, mean accuracy within 80–120%, and mean fragment ion ratio within 70–130% of the coeluting reference fragment ratio (Supplemental Figure 3). Table 2 shows the lowest LLOQ with and without FAIMS from the replicate curves. The full list of LOQs, including ULOQ and range, is shown in Supplemental Table 1. Of 11 precursors, 7 showed a threefold reduction in LLOQ with the addition of FAIMs. For all precursors, the PRM experiments without FAIMS had a median LLOQ of 137 amol/μg; with FAIMS, this was reduced to 46 amol/μg. In parallel, we calculated the limits of quantitation for additional fragment ions (71 in total; ranging from four to ten per precursor), and also for summing all fragment ions signals for each peptide (Supplementary Tables S2 and S3). Summing all fragment ions gave the same result as picking the best fragment ion: the same seven precursors had the lowest LLOQ with FAIMS, while the other four precursors were tied. For the complete set of fragment ions, 37 had the lowest LLOQ with FAIMS, 30 were tied and 4 had the lowest LLOQ without FAIMS. The majority of the ties (22 of 30) were for the same four precursors which did not have an overall improved LLOQ.

**Table 2.**
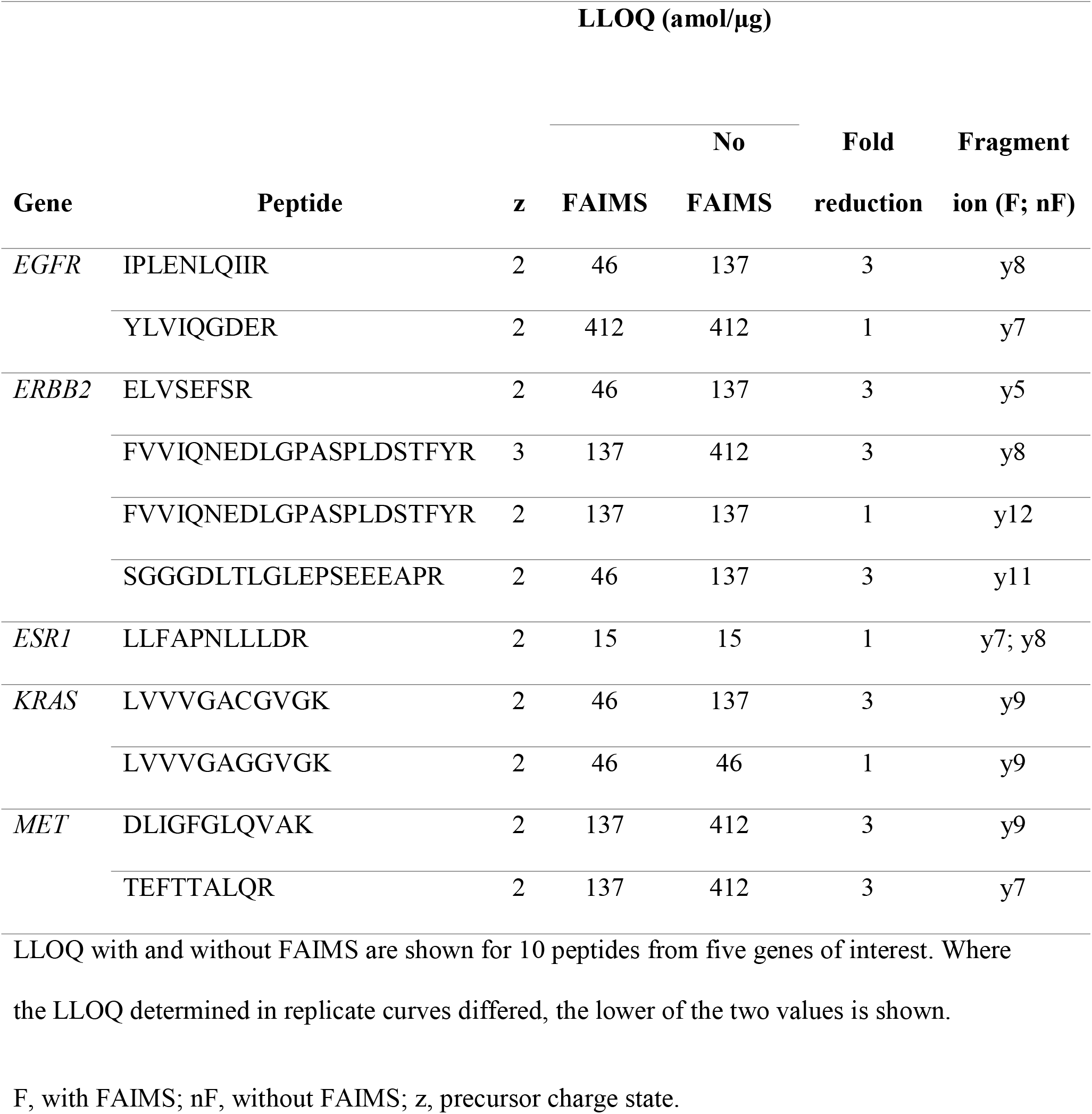
LLOQ comparison

In the case of the EGFR IPLENLQIIR peptide, the lowest LLOQ obtained by PRM analysis without FAIMS was 137 amol (Table 2). This value was improved to 46 amol with FAIMS-PRM. MS/MS spectra of the 46-amol IPLENLQIIR peptide spiked into the spleen matrix and analyzed by FAIMS-PRM vs. PRM are shown in Figure 1C and 1D. At this low level, the PRM MS/MS data showed a higher total ion count (3.9 E5), including interfering background ions, but FAIMS-PRM MS/MS demonstrated a cleaner spectrum, with a fourfold-lower total ion count (9.9 E4) and higher relative intensities of the fragment ions derived from the IPLENLQIIR precursor.

Extracted ion chromatograms (XICs) of fragment ions from the 46-amol IPLENLQIIR peptide analyzed by PRM vs. FAIMS-PRM are shown in Figure 1E and 1F. Interferences detected in PRM, even with high-resolution (30K) Orbitrap detection (Figure 1F), were removed when PRM was acquired with FAIMS (Figure 1E).

We hypothesized that the peptides for which FAIMS reduced the LLOQ in this study were more likely to elute in a region of high matrix complexity with higher levels of non-specific background signal. These peptides would benefit from FAIMS to a greater extent after the background ions were filtered out. The format of our LOQ curve allows us to accurately quantify non-specific background, as this constitutes the only signal in the heavy channel for the LOQ blank injections, with the appropriate elution window defined by the light synthetic peptide. The average number of non-specific background ions injected for each targeted m/z window are shown in Figure 2A. Background ions for all targeted m/z windows decreased with FAIMS (Figure 2B). However, the seven peptides showing a reduction in LLOQ with FAIMS integration demonstrated a significantly greater decrease in background ions with FAIMS. This was shown by a two-sided *t* test comparing percent background reduction for peptides having a lower LLOQ with peptides having an unchanged LLOQ (*P* < 0.001).

**Figure 2.**
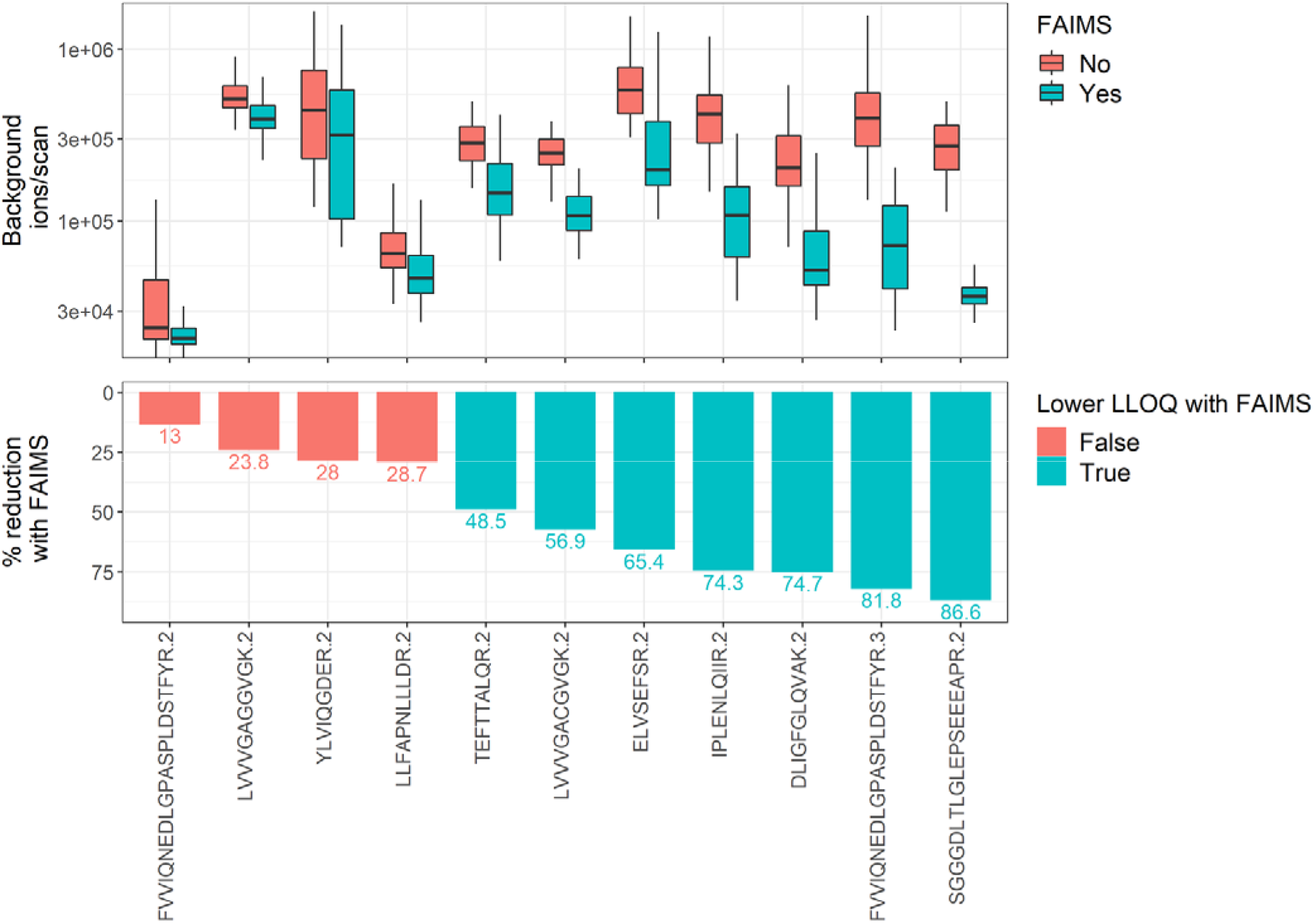
Peptides with a FAIMS-dependent improvement in LLOQ also have a greater reduction in background signal. (A) Background ions per scan, with and without FAIMS for elution time and m/z regions of peptide targets. (B) The percentage reduction in background signal with FAIMS is shown, color-coded by reduction in LLOQ with FAIMS. Background ions are calculated for blank injections and heavy m/z windows only, with the XIC window determined from the light reference peptide elution (n=8, from n=2 LOQ curves per condition; number of scans per plot ranges from 360 to 948). Ion numbers are from ‘RawOvFtT’, extracted from raw files using rawDiag (Trachsel et al., J Proteome Research 2018). Peptides are annotated with precursor sequence and charge-state. Labels on the lower panel indicate the percentage reduction in background signal for each target.

The impact of FAIMS on total ions injected across the LOQ curve is shown in Supplemental Figure S4 for the EGFR IPLENLQIIR [M+2H]^2+^ precursor. The contribution of the spiked heavy peptide was negligible up to 137 amol. From 412 amol, the median ion injection number increased with additional heavy peptide with FAIMS. Without FAIMS, the median ion injection number only noticeably increased from addition of 3.7 fmol heavy peptide, due to the increased background.

### Application of FAIMS-PRM in clinical samples

To demonstrate the performance of our FAIMS-PRM assay on clinical samples, we applied the method to 20 FFPE tumor biopsies from breast cancer patients. Tumor epithelial cells identified by AI-based image analysis were collected by LMD for precise measurement of tumoral concentrations. The overall workflow is described in Figure 3A.

**Figure 3.**
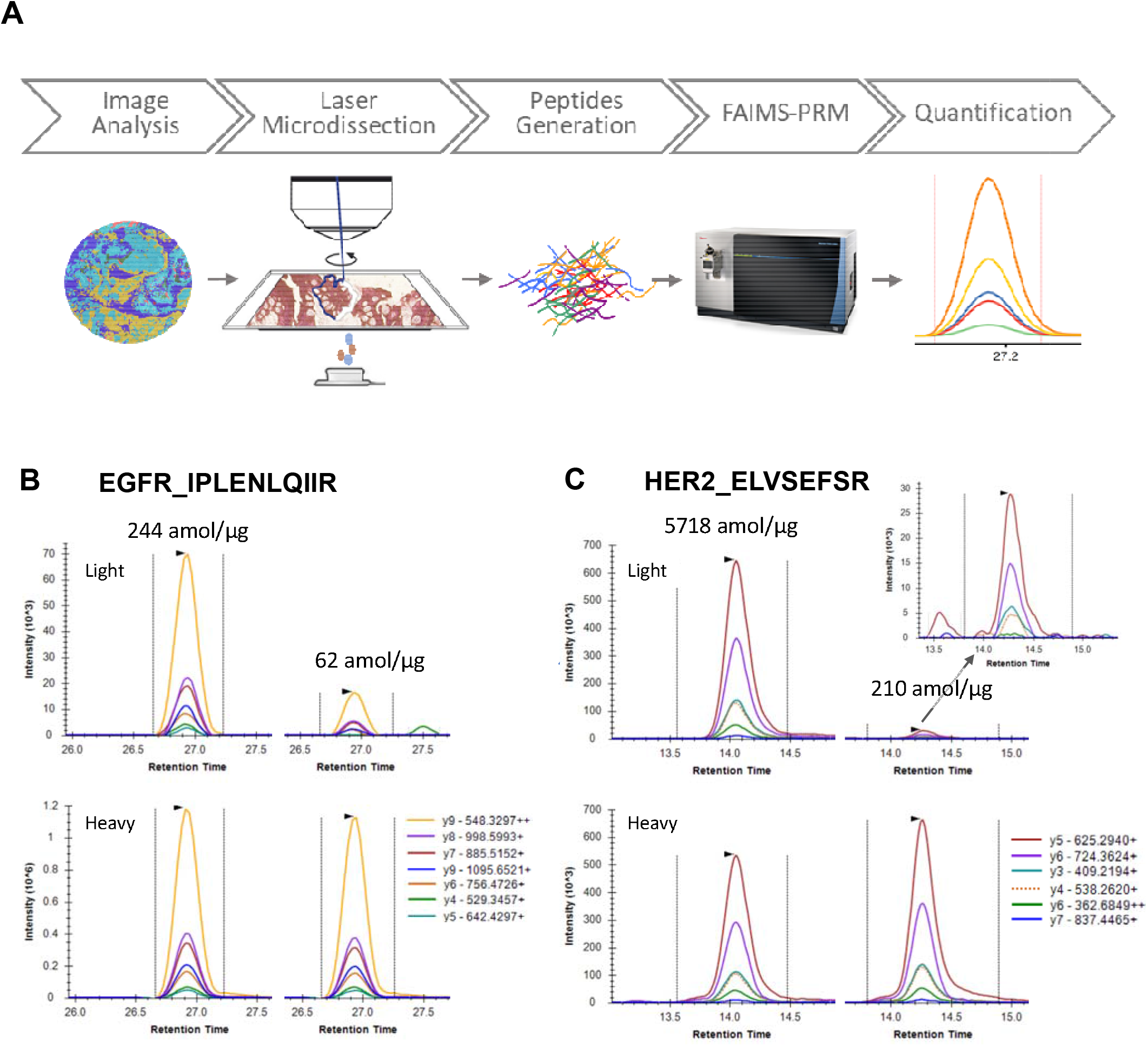
(A) Workflow of clinical proteomics with FAIMS-PRM. (B) Quantitation of EGFR IPLENLQIIR [M+2H]^2+^ from two breast cancer samples. The upper XICs are from endogenous signals, and the lower XICs are from the heavy peptide reference signal (5 fmol). The LLOQ with FAIMS for the y8 fragment ion of this peptide was 46 amol/μg. (C) Quantitation of HER2 ELVSEFSR [M+2H]^2+^ from two breast cancer samples. The upper XICs are from endogenous signal, and the lower XICs are corresponding heavy-peptide reference signals (5 fmol). Inset shows magnified view of the fragment ions for the 210-amol/μg sample. The LLOQ with FAIMS for the y5 fragment ion of this peptide was 46 amol/μg.

All targeted proteins were successfully quantified from the 20 tumor biopsy samples (Supplemental Table S4, Supplemental Figure S5. Examples of FAIMS-PRM–based quantitation for EGFR and HER2 are shown in Figure 3B and 3C. EGFR quantitation from two samples with levels of 244 and 62 amol/μg is shown in Figure 3B. Even at the 62-amol level, the low level of non-specific background signal surrounding the peaks of interest is apparent. Quantitation of HER2 for two samples with levels of 5718 and 210 amol/μg is shown in Figure 3C. HER2 levels in these samples were quantified using the two best-performing peptides, ELVSEFSR and SGGGDLTLGLEPSEEEAPR, which both showed improved LLOQs with FAIMS integration. The HER2 quantitation results obtained independently from these two peptides were highly correlated (R^2^ = 0.991) (Figure 4A).

**Figure 4.**
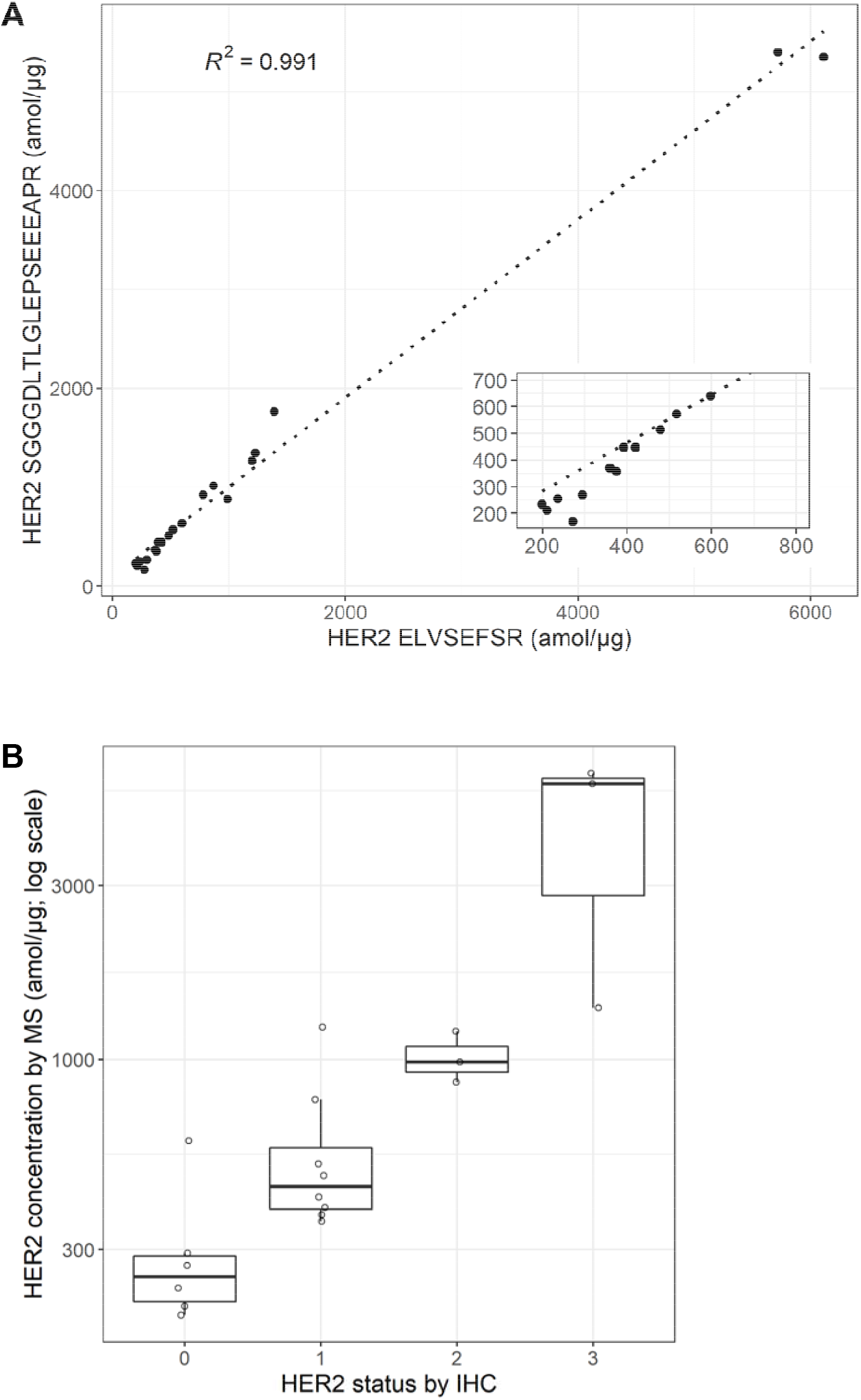
Quantitation of HER2 by FAIMS-PRM in 20 breast cancer biopsies. (A) Scatterplot showing the correlation between HER2 ELVSEFSR and HER2 SGGGDLTLGLEPSEEEAPR peptides. The Pearson correlation coefficient squared value is shown; inset shows expanded view of region below 800 amol/μg. (B) HER2 concentrations by FAIMS-PRM for the ELVSEFSR peptide are shown for each of the HER2 classifications by IHC. All MS values were above the LLOQ.

We also performed IHC analysis for these 20 samples, using the Ventana Her2 (4B5) assay. The pathologist’s scoring results for HER2 expression based on this IHC assay were compared with the tumoral concentrations acquired by FAIMS-PRM. Although the PRM readouts correlated well with the IHC scores, in agreement with earlier publications, the MS quantitation highlighted a wide range of tumoral concentrations within the same IHC scores (Figure 4B). The outliers within an IHC class may identify therapeutic opportunities. E.g. within the eight samples classified as 1+ by IHC, two samples have a HER2 concentration close to the median concentration of the HER2 2+ samples. These patients may be suitable candidates for a HER2-targeted therapy, with a positive FISH test.

## DISCUSSION

Throughout the evolution of targeted proteomics, as in all other analytical methodologies, improved sensitivity has been achieved through limiting noise and increasing signal. The original SRM method continuously transmits ions only within both the precursor and fragment m/z windows, thereby limiting noise from other species and increasing signal ^29^. In practice, multiple transitions per peptide must be monitored to increase confidence in identification and to mitigate potential interference from variable matrix components. However, each additional SRM transition reduces the transmission time per transition and therefore the overall sensitivity of the method. In the PRM method, fragment ion selection is carried out after acquisition from the complete MS/MS data set, and more stringent separation of signal from non-specific background is possible when relying on the high-resolution data. To further increase sensitivity, and if sufficient sample is available (typically at least 10× more than for direct analysis), front-end fractionation or immunoprecipitation can be added into the workflow ^17, 18^. For clinical samples with limited material, incorporation of a gas-phase ion mobility precursor selection step is an alternative approach to enhance sensitivity. We integrated a FAIMS step into the scheduled PRM method to determine whether this integration would improve the quantitation performance for oncology targets in a clinical setting.

The benefit of FAIMS with the EGFR peptide IPLENLQIIR is shown in Figure 1. This peptide showed a dramatic reduction in non-specific background after addition of FAIMS. The effect was evident in the MSMS spectra (Figure 1C vs. 1D) and within the target fragment extracted ion chromatograms (Figure 1E vs. 1F). Lowering the level of non-specific background peptides proximal to the target precursor ion led to detection of more fragment ions with more consistent scan-to-scan signal and consequently higher-quality quantitation.

In the PRM approach with an Orbitrap analyzer, sensitivity can be modulated by ion injection time; longer injection times of up to 500 ms have been shown to improve signal-to-noise ratio by collecting more ions in the C-trap ^30^. As ion injection time is increased, however, throughput is decreased. In our study, we employed a maximum ion injection time of 100 ms per precursor, allowing up to 15 co-eluting peptide pairs to be quantified with six points across a 20-s chromatographic peak width. Dynamic scheduling, based on either elution markers or reference peptide detection, may allow more time to be spent per peptide, further boosting sensitivity ^12, 31^. We note that our 100 ms injection time permits the acquisition of higher resolution data (50K versus 30K), without a significant time penalty. This has been implemented successfully in later version of the method.

The addition of an ion mobility step to an SRM method has been described previously ^19–21^. One report describes a method combining IMS with quadrupole time-of-flight fragment ion detection for the quantitation of host cell proteins in protein biopharmaceutical products ^22^; in this case the IMS was integral to the mass spectrometer, prohibiting a comparison of the same method with and without ion mobility. The FAIMS unit employed in our work has been previously deployed in global proteomic studies, with either label-free or stable-isotope labeling quantitation; to our knowledge, however, its performance in targeted proteomics has not been assessed ^32–36^. Our analysis allowed a direct assessment of the contribution of FAIMS to the performance of the PRM method. Figure 2A shows the variable level of background ions detected within the different precursor windows, quantified as ions per scan, in a human tissue sample. The target of 1E6 ions usually was not reached within the 100-ms fill time. Without FAIMS, the median background level varied by 24-fold, from the highest level of 5.8 E5 ions per scan for the ELVSEFSR (2+) region to the lowest level of 2.4 E4 for the FVVIQNEDLGPASPLDSTFYR (2+) region. With FAIMS, the median background ions per scan was reduced for all 11 precursors, and the degree of reduction ranged from 13 to 86%. The targets with a FAIMS-dependent reduction in LLOQ also showed the greatest FAIMS-dependent reduction in background signal, with a reduction of at least 48.5% (Figure 2B).

We selected five protein targets, HER2, EGFR, cMET, KRAS, and ER, as real-world examples of clinically relevant biomarkers requiring quantitation from small amounts of clinical material—in this case a single 10-μm-thick FFPE section. The tumor areas selected for LMD ranged from 4.6 to 14.1 mm^2^, highlighting the low sample requirements for this sensitive approach. Expression levels for most of these targets varied widely (Supplemental Figure S5). For example, HER2 levels in our samples spanned a 30-fold range, from 200 to 6000 amol/μg (Figure 4). FAIMS-PRM increased the quantitation sensitivity of the clinically relevant biomarker HER2, reducing the LLOQ from 137 to 46 amol/μg. This compares favorably with reported LLOQs from SRM assays of 125 amol/μg ^37^, 5 fmol/μg ^38^, 155 amol/μg ^39^, and 83 amol/μg ^40^, although these assays employed different matrices and LLOQ calculation methods.

Recently, a new anti-HER2 ADC, trastuzumab-deruxtecan (Enhertu; AstraZeneca) showed antitumor activity not only in patients with HER2-positive cancer (HER2 3^+^ and HER2 2^+^/FISH^+^) but also in low-HER2–expressing tumors (HER2 1^+^ and HER2 2^+^) that were previously categorized as HER2 negative and not eligible for anti-HER2 monoclonal antibody treatment ^7^. Importantly, as shown in Figure 4B, HER2 in breast cancer samples still exhibited a range of expression levels in these low-expression samples, and all of them were above the LLOQ of our FAIMS-PRM HER2 assay. Although the tumoral HER2 concentrations used in the box plots are based on the ELVSEFSR peptide, a high correlation (R^2^ = 0.991) was obtained between the assays with two different peptides (ELVSEFSR and SGGGDLTLGLEPSEEEAPR), consistent with the expected quantitation accuracy of our clinical proteomics workflow and FAIMS-PRM assay.

The future development of treatments intended to benefit patients with low HER2 expression will place an increased emphasis on lower-level HER2 quantitation accuracy and robustness. Likewise, the FAIMS-PRM assays for other potential ADC targets such as EGFR and cMET, for which LLOQs were improved (Table 2), will aid investigations of therapeutic benefits for tumors with low target expression. In our study, the addition of FAIMS to the PRM method reduced the LLOQ for most of the peptides evaluated. This increased sensitivity was obtained without additional analysis time or sample material. The clinical proteomics workflow for FFPE tissue samples presented here is anticipated to be of great utility for clinical trials in which multiplexing of protein targets allows proof-of-mechanism and pharmacodynamic questions to be addressed.

## Abbreviations

ADC: antibody-drug conjugate
AI: artificial intelligence
CDx: companion diagnostic
CV: compensation voltage
FAIMS: high-field asymmetric waveform ion mobility spectrometry
FFPE: formalin-fixed, paraffin-embedded
H&E: hematoxylin and eosin
HCD: higher-energy collisional dissociation
IHC: immunohistochemistry
IMS: ion mobility spectrometry
LC: liquid chromatography
LLOQ: lower limit of quantitation
LMS: laser microdissection
LOQ: limit of quantitation
MS: mass spectrometry
MS/MS: tandem mass spectrometry
PRM: parallel reaction monitoring
SRM: selected reaction monitoring
ULOQ: upper limit of quantitation
XIC: extracted ion chromatogram

## Disclosures

### Author contributions

All authors confirmed they have contributed to the intellectual content of this paper and have met the following 4 requirements: (a) significant contributions to the conception and design, acquisition of data, or analysis and interpretation of data; (b) drafting or revising the article for intellectual content; (c) final approval of the published article; and (d) agreement to be accountable for all aspects of the article thus ensuring that questions related to the accuracy or integrity of any part of the article are appropriately investigated and resolved.

## Acknowledgments

We thank Deborah Schuman for proof-reading the manuscript.

## SUPPORTING INFORMATION

Supplemental Table S1. Limits of quantitation for 10 peptides from five proteins of interest; Supplemental Table S2. Summary of lowest LLOQs for summed and individual fragment ions; Supplemental Table S3. Limits of quantitation for summed fragment ions and additional fragment ion quantifiers; Supplemental Table S4. FAIMS-PRM quantification of proteins of interest from 20 breast cancer FFPE tissue samples; Supplemental Figure S1. Increasing isolation width resulted in increased interference and increased signal.; Supplemental Figure S2. Increasing resolution resulted in decreased signal.; Supplemental Figure S3. Illustrative examples of the fragment ratio error calculation; Supplemental Figure S4. Ions injected per scan for EGFR IPLENLQIIR [M+2H]2+, with increasing amounts of spiked-in peptide; Supplemental Figure S5. FAIMS-PRM quantification of proteins of interest from 20 breast cancer samples.; Supplemental Materials and Methods

## Data availability

Skyline files and raw data are available through Panorama, with the ProteomeXchange ID: PXD027134 and access URL: https://panoramaweb.org/0XbJk2.url.

## REFERENCES

1. Vargas, A. J.; Harris, C. C., Biomarker development in the precision medicine era: lung cancer as a case study. Nat Rev Cancer 2016, 16, (8), 525–37.

2. U.S. Food and Drug Administration List of cleared or approved companion diagnostic devices (in vitro and imaging tools), https://www.fda.gov/media/119249/download (March 24, 2021),

3. Scheerens, H.; Malong, A.; Bassett, K.; Boyd, Z.; Gupta, V.; Harris, J.; Mesick, C.; Simnett, S.; Stevens, H.; Gilbert, H.; Risser, P.; Kalamegham, R.; Jordan, J.; Engel, J.; Chen, S.; Essioux, L.; Williams, J. A., Current status of companion and complementary diagnostics: strategic considerations for development and launch. Clin Tronsl Sci 2017, 10, (2), 84–92.

4. Vogel, C. L.; Cobleigh, M. A.; Tripathy, D.; Gutheil, J. C.; Harris, L. N.; Fehrenbacher, L.; Slamon, D. J.; Murphy, M.; Novotny, W. F.; Burchmore, M.; Shak, S.; Stewart, S. J.; Press, M., Efficacy and safety of trastuzumab as a single agent in first-line treatment of HER2-overexpressing metastatic breast cancer. J Clin Oncol 2002, 20, (3), 719–26.

5. Bang, Y. J.; Van Cutsem, E.; Feyereislova, A.; Chung, H. C.; Shen, L.; Sawaki, A.; Lordick, F.; Ohtsu, A.; Omuro, Y.; Satoh, T.; Aprile, G.; Kulikov, E.; Hill, J.; Lehle, M.; Rüschoff, J.; Kang, Y. K., Trastuzumab in combination with chemotherapy versus chemotherapy alone for treatment of HER2-positive advanced gastric or gastro-oesophageal junction cancer (ToGA): a phase 3, open-label, randomised controlled trial. Lancet 2010, 376, (9742), 687–97.

6. An, E.; Ock, C. Y.; Kim, T. Y.; Lee, K. H.; Han, S. W.; Im, S. A.; Kim, T. Y.; Liao, W. L.; Cecchi, F.; Blackler, A.; Thyparambil, S.; Kim, W. H.; Burrows, J.; Hembrough, T.; Catenacci, D. V. T.; Oh, D. Y.; Bang, Y. J., Quantitative proteomic analysis of HER2 expression in the selection of gastric cancer patients for trastuzumab treatment. Ann Oncol 2017, 28, (1), 110–115.

7. Modi, S.; Park, H.; Murthy, R. K.; Iwata, H.; Tamura, K.; Tsurutani, J.; Moreno-Aspitia, A.; Doi, T.; Sagara, Y.; Redfern, C.; Krop, I. E.; Lee, C.; Fujisaki, Y.; Sugihara, M.; Zhang, L.; Shahidi, J.; Takahashi, S., Antitumor activity and safety of trastuzumab deruxtecan in patients with HER2-low–expressing advanced breast cancer: results from a phase Ib study. J Clin Oncol 2020, 38, (17), 1887–1896.

8. Nardone, A.; Weir, H.; Delpuech, O.; Brown, H.; De Angelis, C.; Cataldo, M. L.; Fu, X.; Shea, M. J.; Mitchell, T.; Veeraraghavan, J.; Nagi, C.; Pilling, M.; Rimawi, M. F.; Trivedi, M.; Hilsenbeck, S. G.; Chamness, G. C.; Jeselsohn, R.; Osborne, C. K.; Schiff, R., The oral selective oestrogen receptor degrader (SERD) AZD9496 is comparable to fulvestrant in antagonising ER and circumventing endocrine resistance. Br J Cancer 2019, 120, (3), 331–339.

9. Janes, M. R.; Zhang, J.; Li, L.-S.; Hansen, R.; Peters, U.; Guo, X.; Chen, Y.; Babbar, A.; Firdaus, S. J.; Darjania, L.; Feng, J.; Chen, J. H.; Li, S.; Li, S.; Long, Y. O.; Thach, C.; Liu, Y.; Zarieh, A.; Ely, T.; Kucharski, J. M.; Kessler, L. V.; Wu, T.; Yu, K.; Wang, Y.; Yao, Y.; Deng, X.; Zarrinkar, P. P.; Brehmer, D.; Dhanak, D.; Lorenzi, M. V.; Hu-Lowe, D.; Patricelli, M. P.; Ren, P.; Liu, Y., Targeting KRAS Mutant Cancers with a Covalent G12C-Specific Inhibitor. Cell 2018, 172, (3), 578–589.e17.

10. Nuciforo, P.; Thyparambil, S.; Aura, C.; Garrido-Castro, A.; Vilaro, M.; Peg, V.; Jimenez, J.; Vicario, R.; Cecchi, F.; Hoos, W.; Burrows, J.; Hembrough, T.; Ferreres, J. C.; Perez-Garcia, J.; Arribas, J.; Cortes, J.; Scaltriti, M., High HER2 protein levels correlate with increased survival in breast cancer patients treated with anti-HER2 therapy. Mol Oncol 2016, 10, (1), 138–147.

11. Gallien, S.; Bourmaud, A.; Kim, S. Y.; Domon, B., Technical considerations for large-scale parallel reaction monitoring analysis. J Proteomics 2014, 100, (0), 147–159.

12. Gallien, S.; Kim, S. Y.; Domon, B., Large-scale targeted proteomics using internal standard triggered-parallel reaction monitoring (IS-PRM). Mol Cell Proteomics 2015, 14, (6), 1630–1644.

13. Gallien, S.; Duriez, E.; Crone, C.; Kellmann, M.; Moehring, T.; Domon, B., Targeted proteomic quantification on quadrupole-Orbitrap mass spectrometer. Mol Cell Proteomics 2012, 11, (12), 1709–1723.

14. Cifani, P.; Kentsis, A., High sensitivity quantitative proteomics using automated multidimensional nano-flow chromatography and accumulated ion monitoring on quadrupole-Orbitrap-linear ion trap mass spectrometer. Mol Cell Proteomics 2017, 16, (11), 2006–2016.

15. Kim, Y. J.; Gallien, S.; van Oostrum, J.; Domon, B., Targeted proteomics strategy applied to biomarker evaluation. Proteomics Clin Appl 2013, 7, (11-12), 739–47.

16. Peterson, A. C.; Russell, J. D.; Bailey, D. J.; Westphall, M. S.; Coon, J. J., Parallel reaction monitoring for high resolution and high mass accuracy quantitative, targeted proteomics. Mol Cell Proteomics 2012, 11, (11), 1475–88.

17. Morales-Betanzos, C. A.; Lee, H.; Gonzalez Ericsson, P. I.; Balko, J. M.; Johnson, D. B.; Zimmerman, L. J.; Liebler, D. C., Quantitative mass spectrometry analysis of PD-L1 protein expression, *N*-glycosylation and expression stoichiometry with PD-1 and PD-L2 in human melanoma. Mol Cell Proteomics 2017, 16, (10), 1705–1717.

18. Do, M.; Kim, H.; Yeo, I.; Lee, J.; Park, I. A.; Ryu, H. S.; Kim, Y., Clinical application of multiple reaction monitoring-mass spectrometry to human epidermal growth factor receptor 2 measurements as a potential diagnostic tool for breast cancer therapy. Clin Chem 2020, 66, (10), 1339–1348.

19. Klaassen, T.; Szwandt, S.; Kapron, J. T.; Roemer, A., Validated quantitation method for a peptide in rat serum using liquid chromatography/high-field asymmetric waveform ion mobility spectrometry. Rapid Commun Mass Spectrom 2009, 23, (15), 2301–2306.

20. Xia, Y.-Q.; Wu, S. T.; Jemal, M., LC-FAIMS-MS/MS for quantification of a peptide in plasma and evaluation of FAIMS global selectivity from plasma components. Anal Chem 2008, 80, (18), 7137–7143.

21. Norman, R. L.; Singh, R.; Langridge, J. I.; Ng, L. L.; Jones, D. J. L., The measurement of KRAS G12 mutants using multiplexed selected reaction monitoring and ion mobility mass spectrometry. Rapid Commun Mass Spectrom 2020, 34, (S4), e8657.

22. Doneanu, C.; Fang, J.; Alelyunas, Y.; Yu, Y. Q.; Wrona, M.; Chen, W., An HS-MRM assay for the quantification of host-cell proteins in protein biopharmaceuticals by liquid chromatography ion mobility QTOF mass spectrometry. J Vis Exp 2018, (134), e55325.

23. Cooper, H. J., To what extent is FAIMS beneficial in the analysis of proteins? J Am Soc Mass Spectrom 2016, 27, (4), 566–577.

24. Renuse, S.; Vanderboom, P. M.; Maus, A. D.; Kemp, J. V.; Gurtner, K. M.; Madugundu, A. K.; Chavan, S.; Peterson, J. A.; Madden, B. J.; Mangalaparthi, K. K.; Mun, D.-G.; Singh, S.; Kipp, B. R.; Dasari, S.; Singh, R. J.; Grebe, S. K.; Pandey, A., A mass spectrometry-based targeted assay for detection of SARS-CoV-2 antigen from clinical specimens. EBioMedicine 2021, 69, 103465.

25. MacLean, B.; Tomazela, D. M.; Shulman, N.; Chambers, M.; Finney, G. L.; Frewen, B.; Kern, R.; Tabb, D. L.; Liebler, D. C.; MacCoss, M. J., Skyline: an open source document editor for creating and analyzing targeted proteomics experiments. Bioinformatics 2010, 26, (7), 966–968.

26. Trachsel, C.; Panse, C.; Kockmann, T.; Wolski, W. E.; Grossmann, J.; Schlapbach, R., rawDiag: an R package supporting rational LC–MS method optimization for bottom-up proteomics. J Proteome Res 2018, 17, (8), 2908–2914.

27. Sellmann, C.; Doerner, A.; Knuehl, C.; Rasche, N.; Sood, V.; Krah, S.; Rhiel, L.; Messemer, A.; Wesolowski, J.; Schuette, M.; Becker, S.; Toleikis, L.; Kolmar, H.; Hock, B., Balancing selectivity and efficacy of bispecific epidermal growth factor receptor (EGFR) × c-MET antibodies and antibody-drug conjugates. J Biol Chem 2016, 291, (48), 25106–25119.

28. Perry, R. H.; Cooks, R. G.; Noll, R. J., Orbitrap mass spectrometry: instrumentation, ion motion and applications. Mass Spectrom Rev 2008, 27, (6), 661–99.

29. Picotti, P.; Aebersold, R., Selected reaction monitoring–based proteomics: workflows, potential, pitfalls and future directions. Nat Methods 2012, 9, 555–566.

30. Nguyen, C. D. L.; Malchow, S.; Reich, S.; Steltgens, S.; Shuvaev, K. V.; Loroch, S.; Lorenz, C.; Sickmann, A.; Knobbe-Thomsen, C. B.; Tews, B.; Medenbach, J.; Ahrends, R., A sensitive and simple targeted proteomics approach to quantify transcription factor and membrane proteins of the unfolded protein response pathway in glioblastoma cells. Sci Rep 2019, 9, (1), 8836.

31. Remes, P. M.; Yip, P.; MacCoss, M. J., Highly multiplex targeted proteomics enabled by real-time chromatographic alignment. Anal Chem 2020, 92, (17), 11809–11817.

32. Pfammatter, S.; Bonneil, E.; McManus, F. P.; Prasad, S.; Bailey, D. J.; Belford, M.; Dunyach, J.-J.; Thibault, P., A novel differential ion mobility device expands the depth of proteome coverage and the sensitivity of multiplex proteomic measurements. Mol Cell Proteomics 2018, 17, (10), 2051–2067.

33. Bekker-Jensen, D. B.; del Val, A. M.; Steigerwald, S.; Rüther, P.; Fort, K.; Arrey, T. N.; Harder, A.; Makarov, A.; Olsen, J. V., A compact quadrupole-Orbitrap mass spectrometer with FAIMS interface improves proteome coverage in short LC gradients. Preprint at http://biorxiv.org/content/early/2019/11/30/860643.abstract.

34. Hebert, A. S.; Prasad, S.; Belford, M. W.; Bailey, D. J.; McAlister, G. C.; Abbatiello, S. E.; Huguet, R.; Wouters, E. R.; Dunyach, J.-J.; Brademan, D. R.; Westphall, M. S.; Coon, J. J., Comprehensive singleshot proteomics with FAIMS on a hybrid Orbitrap mass spectrometer. Anal Chem 2018, 90, (15), 9529–9537.

35. Pfammatter, S.; Bonneil, E.; McManus, F. P.; Thibault, P., Accurate quantitative proteomic analyses using metabolic labeling and high field asymmetric waveform ion mobility spectrometry (FAIMS). J Proteome Res 2019, 18, (5), 2129–2138.

36. Schweppe, D. K.; Prasad, S.; Belford, M. W.; Navarrete-Perea, J.; Bailey, D. J.; Huguet, R.; Jedrychowski, M. P.; Rad, R.; McAlister, G.; Abbatiello, S. E.; Woulters, E. R.; Zabrouskov, V.; Dunyach, J.-J.; Paulo, J. A.; Gygi, S. P., Characterization and optimization of multiplexed quantitative analyses using high-field asymmetric-waveform ion mobility mass spectrometry. Anal Chem 2019, 91, (6), 4010–4016.

37. Hembrough, T.; Thyparambil, S.; Liao, W.-L.; Darfler, M. M.; Abdo, J.; Bengali, K. M.; Hewitt, S. M.; Bender, R. A.; Krizman, D. B.; Burrows, J., Application of selected reaction monitoring for multiplex quantification of clinically validated biomarkers in formalin-fixed, paraffin-embedded tumor tissue. J Mol Diagn 2013, 15, (4), 454–465.

38. Guerin, M.; Gonçalves, A.; Toiron, Y.; Baudelet, E.; Pophillat, M.; Granjeaud, S.; Fourquet, P.; Jacot, W.; Tarpin, C.; Sabatier, R.; Agavnian, E.; Finetti, P.; Adelaide, J.; Birnbaum, D.; Ginestier, C.; Charafe-Jauffret, E.; Viens, P.; Bertucci, F.; Borg, J. P.; Camoin, L., Development of parallel reaction monitoring (PRM)-based quantitative proteomics applied to HER2-Positive breast cancer. Oncotarget 2018, 9, (73), 33762–33777.

39. Steiner, C.; Tille, J.-C.; Lamerz, J.; Kux van Geijtenbeek, S.; McKee, T. A.; Venturi, M.; Rubbia-Brandt, L.; Hochstrasser, D.; Cutler, P.; Lescuyer, P.; Ducret, A., Quantification of HER2 by targeted mass spectrometry in formalin-fixed paraffin-embedded (FFPE) breast cancer tissues. Mol Cell Proteomics 2015, 14, (10), 2786–2799.

40. Kennedy, J. J.; Abbatiello, S. E.; Kim, K.; Yan, P.; Whiteaker, J. R.; Lin, C.; Kim, J. S.; Zhang, Y.; Wang, X.; Ivey, R. G.; Zhao, L.; Min, H.; Lee, Y.; Yu, M.-H.; Yang, E. G.; Lee, C.; Wang, P.; Rodriguez, H.; Kim, Y.; Carr, S. A.; Paulovich, A. G., Demonstrating the feasibility of large-scale development of standardized assays to quantify human proteins. Nat Methods 2013, 11, 149.

