## Supplemental Data for "The addition of FAIMS Increases Targeted Proteomics Sensitivity from FFPE Tumor Biopsies"

AstraZeneca, Gaithersburg, MD

Table of Contents

**Supplemental Figure S1.** Increasing isolation width resulted in increased interference and increased signal.	S-2

**Supplemental Figure S2.** Increasing resolution resulted in decreased signal. .	S-4

**Supplemental Figure S3.** Illustrative examples of the fragment ratio error calculation.	S-5

**Supplemental Figure S4.** Ions injected per scan for EGFR IPLENLQIIR [M+2H]2+, with increasing amounts of spiked-in peptide..	S-6

**Supplemental Figure S5.** Quantification of proteins of interest from 20 breast cancer samples.	S-7

**Supplemental Materials and Methods.**.	S-8

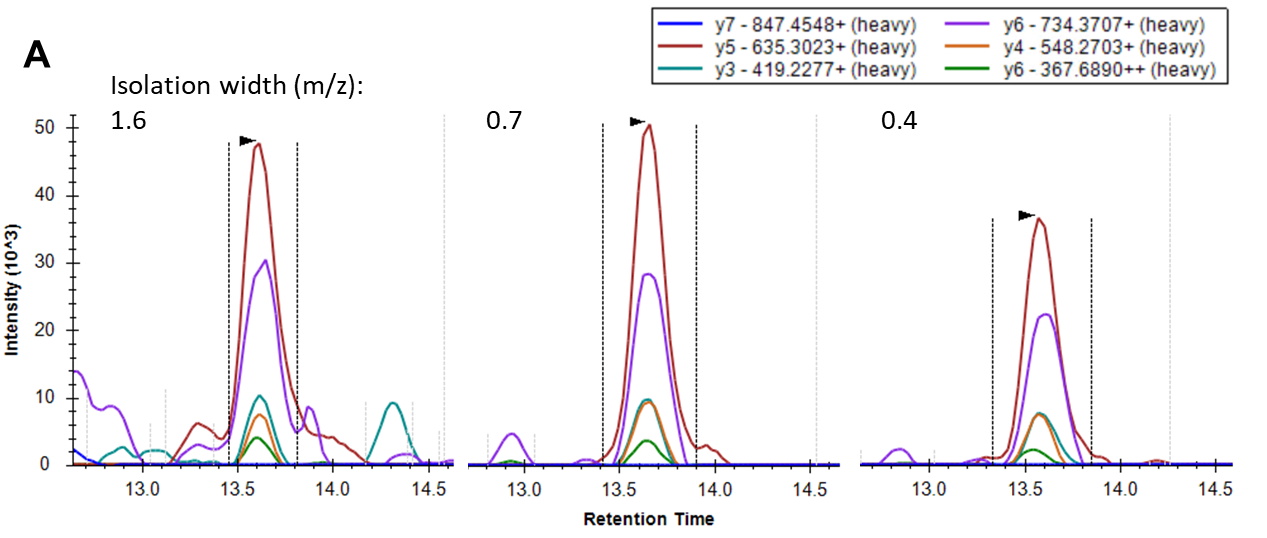

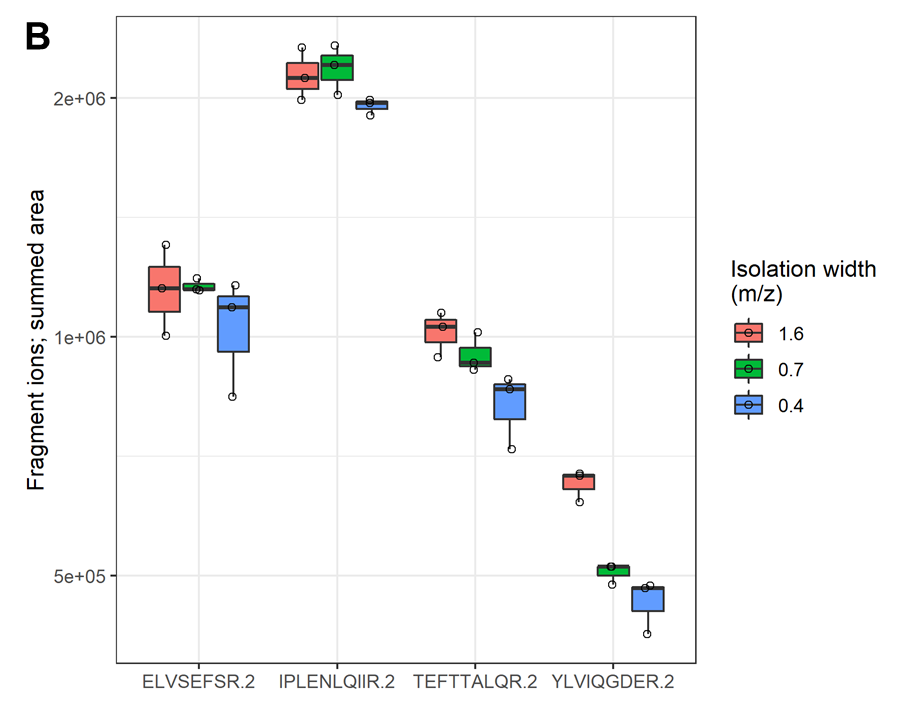

**Supplemental Figure S1.** Increasing isolation width resulted in increased interference and increased signal. A) Extracted ion chromatograms of FAIMS-PRM fragment ions of heavy HER2 ELVSEFSR [M+2H]^2+^ at 200 amol in 1 μg of spleen matrix acquired with isolation widths of 1.6, 0.7 and 0.4 m/z. B) The summed fragment ion area for 200 amol heavy peptide in 1 µg spleen matrix is shown for each isolation width, acquired in triplicate. The resolution was constant, at 30K.

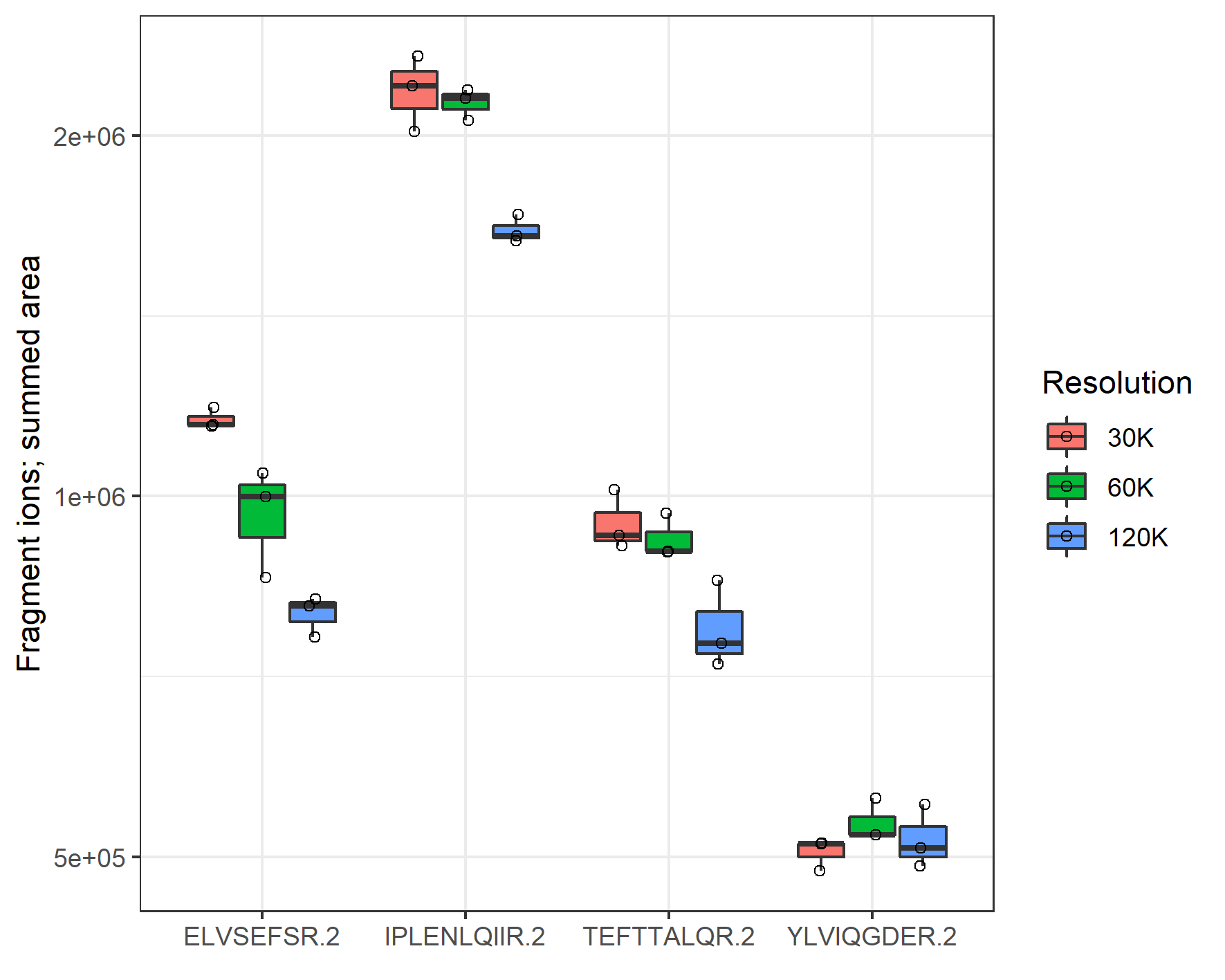

**Supplemental Figure S2.** Increasing resolution resulted in decreased signal. The summed fragment ion area for 200 amol heavy peptide in 1 µg spleen matrix is shown for FAIMS-PRM of four target peptides at each resolution, acquired in triplicate. The isolation width was constant, at 0.7 m/z.

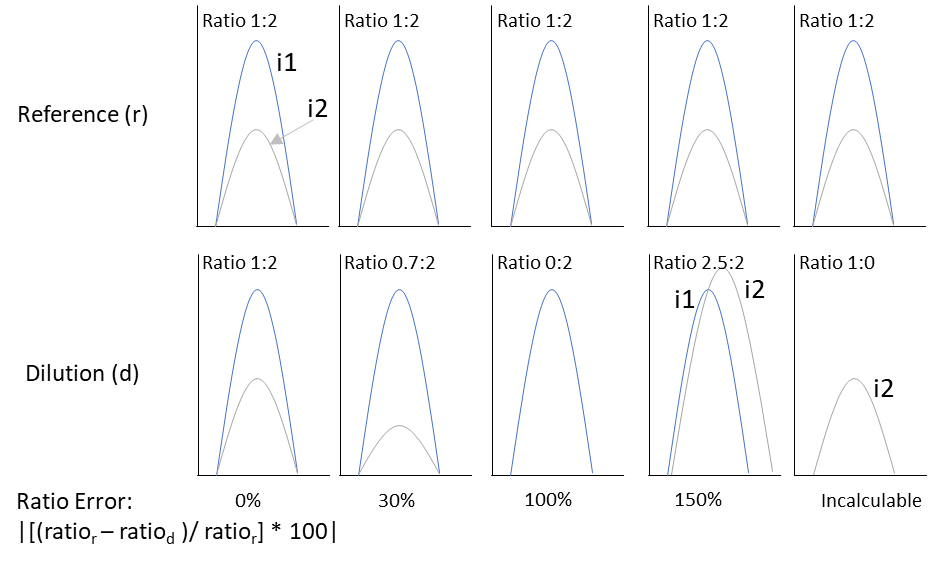

**Supplemental Figure S3.** Illustrative examples of the fragment ratio error calculation. The i1 (blue) and i2 (grey) fragment ions are defined from the reference intensities. From left to right the cartoons illustrate: 1) perfect agreement between reference and sample/dilution ratios; 2) high i1 relative intensity in the sample/dilution data, e.g. due to interference; 3) missing i2 fragment ion in the sample/dilution data; 4) high i2 relative intensity in the sample/dilution data, e.g. due to interference; 5) missing i1 fragment ion in the sample/dilution data.

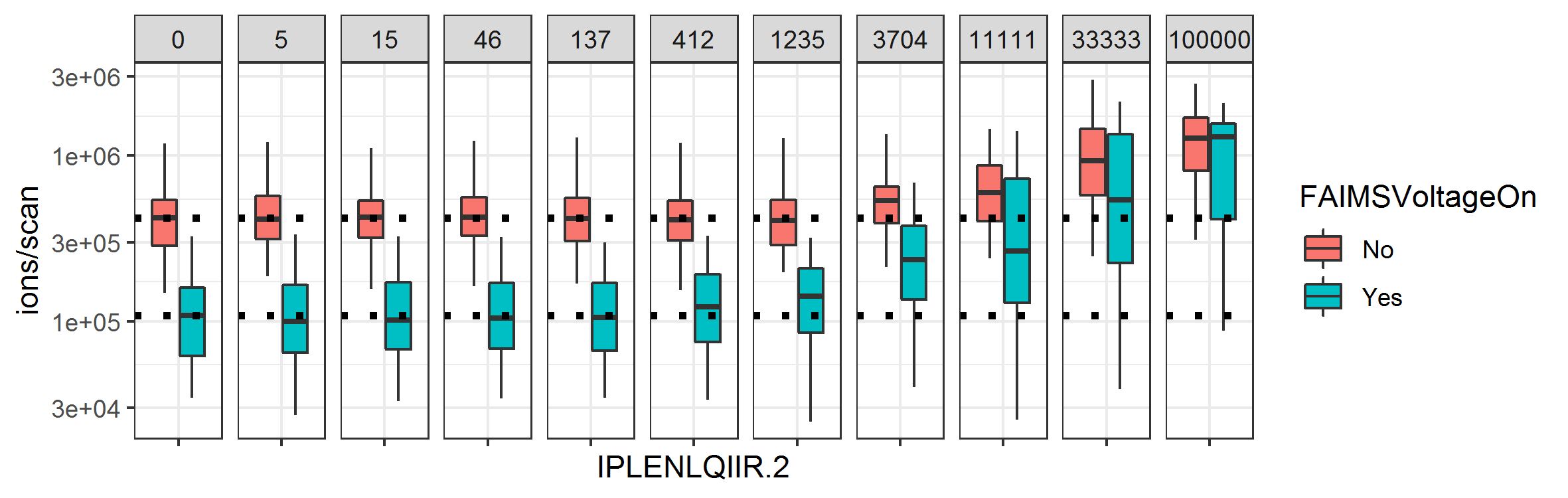

FAIMS

Heavy peptide spiked in (amol)

**Supplemental Figure S4**. Ions injected per scan for EGFR IPLENLQIIR [M+2H]^2+^, with increasing amounts of spiked-in peptide. From 0 (non-specific background only) to 100 fmol (mainly specific signal); heavy m/z window only. Horizontal dotted lines indicate the median values for no heavy peptide.

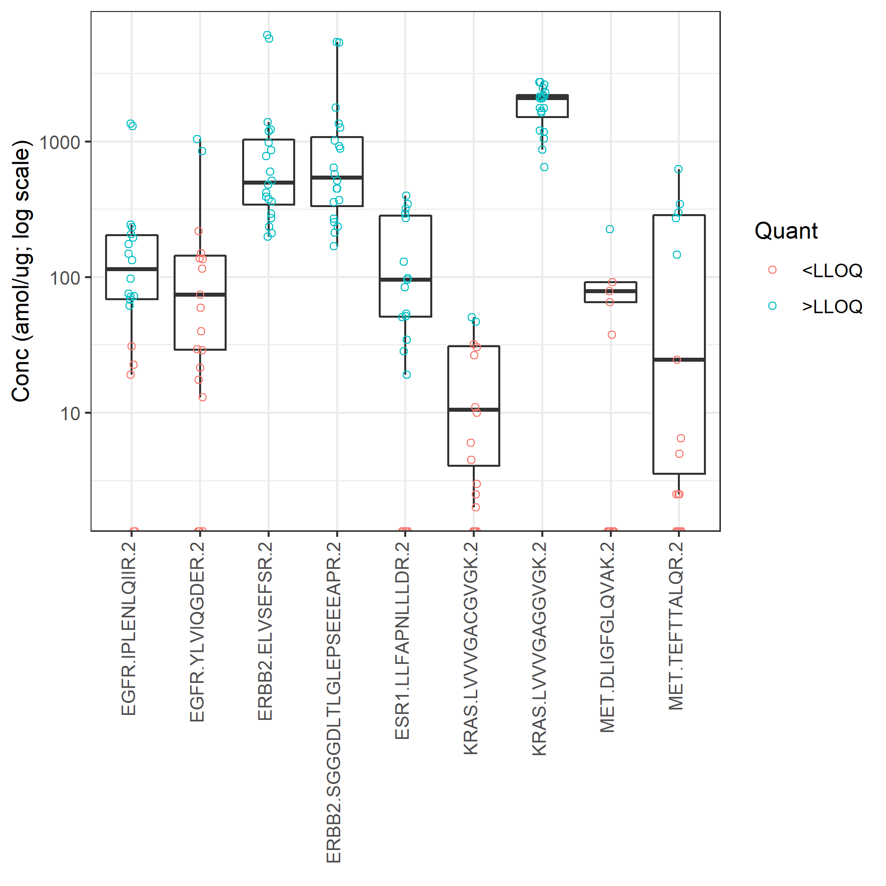

**Supplemental Figure S5.** FAIMS-PRM quantification of proteins of interest from 20 breast cancer samples. Each data point represents a clinical sample. Y-axis is log-scale; 44 data points with quantitation of 0 are omitted from the plot.

**Supplemental Materials and Methods**

**Image analysis and LMD**

Pathology evaluation for laser-microdissected tumor epithelium was conducted by the study pathologist on H&E-stained slides, which were digitally imaged with an Aperio ScanScope AT scanner (Leica Microsystems, Wetzlar, Germany). HALO AI (Indica Labs, Albuquerque, NM) was used to classify and annotate the slides to guide LMD of tumor epithelium. The HALO AI minNet classification tool was used with training annotations: *n* = 199 for tumor epithelium, *n* = 148 stroma, *n* = 44 inflammatory infiltrates, *n* = 23 necrosis, and *n* = 19 glass/empty space/red blood cells/tissue artifacts.

FFPE tissue sections adjacent to the H&E slide were sectioned at a 10-µm thickness and mounted onto UV-treated polyethylene membrane slides (Leica). The tissue sections were deparaffinized with two changes in xylene and hydrated through descending grades of ethanol (100%, 95%, and 75%) and deionized water. The slides were stained with hematoxylin (Gill 2; Richard Allan Scientific, San Diego, CA) and subsequently dehydrated through deionized water and ascending grades of ethanol (75%, 95%, and 100%) and xylene. Hematoxylin-stained slides were allowed to air dry for at least 5 min before LMD, which was performed with the LMD 6500 Laser Microdissection System (Leica) and a dissection time maintained at under 1 h per slide. Based on the HALO AI image markups for tumor epithelium, slide annotations were hand drawn, using the 5× objective. Annotated targets were cut with a UV laser, using the 5× objective with the laser screw setting, advanced mode (step size, 2 µm; repeats, 3). The following laser parameters were used: laser power, 51%; aperture, 29; speed, 32; specimen balance, 6. Dissected targets were collected in the cap of a 0.5-mL microcentrifuge tube (Applied Biosystems, Foster City, CA). Representative images were then documented for each sample and included the target area both before LMD and after laser capture, the LMD area after target removal, and the collection cap with target material. Material was suspended in 30 µL of acetonitrile, and the cap containing the sample was placed onto an Eppendorf LoBind 0.5-mL microcentrifuge tube (Thermo Fisher Scientific, Waltham, MA). Sample was transferred by microcentrifugation at 13,300 rpm for 2 min and stored at –30°C.

**Sample processing for LC-MS analysis**

Microdissected tissue samples collected from the slides were dissolved in 0.1% RapiGest (186001861; Waters, Milford, MA) and 100 mM tris(hydroxymethyl)aminomethane (T2694; Sigma Chemical Company, St. Louis, MO) before incubation at 95°C with shaking at 1,000 rpm for 90 min in an Eppendorf ThermoMixer C (Sigma). Tris(2-carboxyethyl)phosphine (77720; Thermo Fisher) and chloroacetamide (C0267; Sigma) were added to final concentrations of 10 and 34 mM, respectively, and samples were incubated at 37°C, 500 rpm for 30 min. Trypsin (V5111; Promega, Madison, WI) was added, and incubation continued at 37°C at 500 rpm for 16–18 h. Samples were subsequently acidified with trifluoroacetic acid at a final concentration of 1% and incubated at 37°C, 500 rpm for 30 min. Samples were then centrifuged at 17,000 RCF for 10 min. The supernatant was desalted by using a reversed-phase sorbent (Oasis HLB 96-well µElution Plate, 186001828BA; Waters), with elution in 70 µL of 50% acetonitrile with 0.1% formic acid. The samples were dried by vacuum centrifugation and subsequently resuspended in aqueous 0.1% formic acid with shaking at 50°C, 1,000 rpm for 10 min. Peptide concentration was determined using a bicinchoninic acid assay (Micro BCA Protein Assay Kit, 23235; Thermo Fisher) with 10% radioimmunoprecipitation buffer as diluent (89900; Pierce Chemical, Dallas, TX).

Bulk formalin-fixed human spleen, homogenized and prepared as described above, was used as a complex matrix to determine the limit of quantitation (LOQ). A reverse-dilution LOQ experiment was carried out with spiking of constant light synthetic peptide (5 fmol) and a dilution series of heavy stable-isotope–labeled synthetic peptide. A 10-point dilution series was acquired in quadruplicate, ranging from 5 amol to 100 fmol on-column. Additional single and double blanks (reference peptide plus matrix; matrix only) were also acquired in quadruplicate.

**LC-MS data acquisition**

Samples (1 µg, spiked with 5 fmol isotope-labelled synthetic peptides) were loaded onto EvoTip trapping columns before separation with the EvoSep One nanoLC system (EvoSep, Odense, Denmark) coupled to an Orbitrap Fusion Lumos mass spectrometer with a FAIMS-PRO interface (Thermo Fisher). Peptides were eluted over a 44-min gradient, from 7% to 30% acetonitrile (on-column), at a flow rate of 500 nL/min. The mobile phase comprised buffer A, 0.1% aqueous formic acid (LS118-4; Thermo Fisher), and mobile phase B comprised 0.1% formic acid in acetonitrile (LS120-212; Thermo Fisher). The analytical column was 15 cm × 100 µm with 3-µm particles (15-100-3-UHPnC; PepSep, Marslev, Denmark). A stainless steel emitter was employed (30 × 150 µm, PSSELJ; PepSep) with an applied voltage of 2.4 kV. The FAIMS-PRM experiment employed higher-energy collisional dissociation (HCD) fragmentation with an isolation window of 0.7 mass-to-charge ratio (m/z), a target automatic gain control of 1E6 ions, and a maximum injection time of 100 ms. Tandem MS (MS/MS) scans were acquired in centroid mode with the Orbitrap detector, using 30K resolution at 200 m/z. FAIMS was operated at the standard resolution, with no additional FAIMS gas.

**Data analysis**

PRM data was analyzed with Skyline (University of Washington, Seattle, WA) (24), using high-selectivity extraction; for a given m/z value, this extraction window corresponds to:
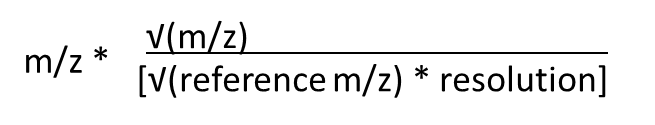
. Fragment ions with interference were identified by manual analysis, comparing coelution and fragment ion ratios between endogenous and reference peptides. Any fragment ions showing interference were flagged and omitted from use in quantitation. Quantitation was performed from a single fragment ion, with the amount calculated from the fragment “Area Ratio” exported from Skyline and multiplied by the internal standard amount (5 fmol).

A 10-point standard curve spanning 5–100 fmol was used to determine the lower LOQ (LLOQ) and upper LOQ (ULOQ). A linear regression with 1/x (analyte amount) weighting was carried out in R with the linear model function, using fragment ion peak area ratios exported from Skyline. The measured amount was extrapolated from the linear regression for each point. Mean fragment ion accuracy was required to be within 80–120% of the expected value for all points from the LLOQ to the ULOQ. CV was required to be less than 20% for all points from the LLOQ to the ULOQ. The mean fragment ion ratio for the top two quantitative fragments (i.e., not flagged as interfering) was required to be within 30% of that for the coeluting standard fragment for all points from the LLOQ to the ULOQ. The fragment ion ratio error was calculated as follows: (1) the first- and second-most intense fragments (i1 and i2) were defined from the coeluting reference data, (2) the ratio for i2:i1 was calculated for reference and curve data, and (3) the error was calculated as the absolute value of [(curve fragment ratio – reference fragment ratio)/reference fragment ratio] *100. For example, with a reference fragment ratio of 1:2 and a curve fragment ratio of 1:4, we obtained abs{[(0.25 – 0.5)/0.5] *100} = 50%. For an extreme cases, if the curve first-intensity fragment was 0 (1:0), the ratio and ratio error was infinity; if both the curve first- and second-intensity fragments were missing, the ratio and ratio error were incalculable (NA); if the curve second-intensity fragment was 0, the ratio was 0 and the ratio error was 100%. The single-fragment ion with the lowest LLOQ was selected for each peptide. Where multiple fragments had the same LLOQ, the fragment with the higher ULOQ was preferred. Where the LLOQ and ULOQ were identical, the more intense fragment was preferred.
